# Maternal dietary deficiencies in folic acid and choline change metabolites levels in offspring after ischemic stroke

**DOI:** 10.1101/2024.06.29.601358

**Authors:** Fazian Anwar, Mary-Tyler Mosley, Paniz Jasbi, Jinhua Chi, Haiwei Gu, Nafisa M. Jadavji

## Abstract

Ischemic stroke is a debilitating disease, with nutrition being a modifiable risk factor. Changes in levels of metabolites can be used to measure the alterations in the gut, a significant marker for the etiology of diseases. This study utilized untargeted metabolomics to investigate changes in fecal samples of offspring in response to maternal dietary deficiencies and ischemic stroke. Female mice were placed on control (CD), folic acid- (FADD), or choline-deficient (ChDD) diets prior to, during pregnancy, and lactation. Offspring were weaned on to CD and at 2 months of age an ischemic stroke was induced. Fecal samples were collected prior to ischemic stroke, and at 1- and 4-weeks post-stroke for analysis. Sex and maternal dietary differences in metabolites were observed at both the 1- and 4-week post-stroke timepoints. At the 1-week post-stroke, female FADD offspring had more changes in metabolites than males. Comparatively, at the 4-week post-stroke timepoint, female offspring on either FADD or ChDD demonstrated metabolite changes. This study demonstrates a long-lasting impact of maternal dietary deficiencies on central nervous system and gut microbiome function after ischemic stroke.

**Summary Statement:** Our study investigated metabolite changes in female and male offspring fecal samples from mothers maintained on folic acid or choline deficient diet. We report that female offspring metabolite levels were impacted.

## Introduction

Ischemic stroke is one of the leading causes of morbidity and mortality globally. The incidence of stroke in adults has been increasing due to factors such as high blood pressure, vascular disorders, diabetes, and obesity (Mozaffarian et al., 2015). By 2030, the number of individuals affected by stroke is predicted to increase by 3.4 million (Ovbiagele et al., 2013). While there has been a higher prevalence of ischemic stroke in older age individuals (> 65 years) (Feigin et al., 2014; Mozaffarian et al., 2015), recent studies have shown that the incidence of stroke among people under 50 years of age is alarmingly increasing (Feigin et al., 2014; Smajlović, 2015; Tong et al., 2016). More importantly it is becoming more evident that nutrition is a well-established and a modifiable risk factor for vascular dysfunction leading to ischemic stroke (Benjamin et al., 2018; Hankey, 2012; Spence et al., 2017).

Although previous studies have established nutrition as a modifiable risk factor for vascular dysfunction and ischemic stroke (Benjamin et al., 2018; Hankey, 2012; Spence et al., 2017), logistical challenges in conducting longitudinal studies have limited research on the impact of early-life nutrition on late-life neurological outcomes. Pregnancy and early life are critical periods for neurodevelopment and lay the foundation for future brain function and health (Carolan-Olah et al., 2015; Kolb et al., 2016; Pannia et al., 2016). In the US, health disparities lead many women to receive inadequate vitamins and nutrients during pregnancy. For example, nearly one in three African-American women do not get enough folic acid daily (Bailey et al., 2010). The same is true for Spanish-speaking Mexican-American women; they often do not get enough folic acid (Hamner et al., 2011). Furthermore, approximately one-fourth of women in the US do not consume enough choline from their diet (Jensen et al., 2007; Zeisel, 2013), and women in low-income countries often face nutritional deficiencies during pregnancy, making this a serious global health issue (Gernand et al., 2016). Despite recent attention to the potential effects of maternal dietary deficiency on early life and childhood development, the long-term impact of maternal dietary deficiencies during pregnancy on the child’s future vascular health is not yet well understood. Evidence suggests that maternal nutrition plays an important role in disease onset and outcome later in life (Di Meco & Praticò, 2019; Gaillard, 2015; Palinski, 2014; Yessoufou & Moutairou, 2011). For example, maternal obesity during pregnancy is implicated in the onset of metabolic diseases, such as diabetes in offspring( Gaillard, 2015; Yessoufou and Moutairou, 2011), but more life course studies are required (Gaillard, 2015). Furthermore, it has been shown that the maternal environment plays an important role in cardiovascular risk in offspring (Palinski, 2014). Using animal models, many groups have shown that maternal diet (e.g., high-fat diet, and caloric restriction, etc.) can modulate cerebrovascular structure and function, leading to endothelial dysfunction, reduced smooth muscle contractility, and decreased myogenic tone, all of which significantly influence peripheral and cerebral vascular health in mouse and rat adult offspring (Campbell et al., 2005; Durrant et al., 2014; Lin et al., 2018). Our group has demonstrated that maternal dietary deficiencies in folic acid or choline impact offspring ischemic stroke outcome (Clementson et al., 2023; Hurley et al., 2023; Pull et al., 2023). However, the mechanisms are not yet clear, our hypothesis is that maternal diet deficient in either folic acid or choline combined with ischemic stroke will result in changes to metabolites within the gut. The aim of the present study was to investigate changes in metabolite levels of fecal matter in male and female offspring exposed to FADD and ChDD during early neurodevelopment and an ischemic stroke during adulthood. Untargeted metabolite analysis has been shown to provide some insight to changes after stroke (Zhao et al., 2022), predict stroke outcome (Chi et al., 2021), as well as generate data for biomarkers (Li et al., 2018; M.-H. Wu et al., 2023).

## Results

Maternal diet deficiencies in FADD or ChDD during pregnancy and lactation impacted metabolite levels at the 1- and 4-week post-stroke timepoints. The number of changes in metabolites because of experimental manipulations (maternal diet or offspring sex) are summarized in Figure 1. The number of metabolites measured in offspring fecal samples was extensive. We have listed all the metabolites that were significantly difference in supplementary Table 1 with the corresponding p-values for interactions between maternal diet and offspring sex, as well as main effects of maternal diet and sex.

**Figure 1.** Summary of two-way ANOVA analysis demonstrating the metabolite changes based on sex, maternal diet, or interaction. (A) Demonstrate difference in one metabolite pre-stroke due to change in maternal diet. (B) Demonstrate metabolite changes based on sex, maternal diet, and combined effect of both maternal diet & sex in 1-week post-stroke with no interaction. (C) Significant and enhanced changes in metabolites seen at 4-weeks post stroke based on sex, maternal diet, or combined with 3 metabolites changes being affected differently by sex and maternal diet.

### Pre-Stroke

At the pre-stroke time point, a significant difference in HEPES levels was observed between maternal diet groups (p = 0.00052), with female ChDD offspring having elevated levels of this metabolite (Table 1). There was also a difference between male and female offspring (p = 0.0087) and interaction between maternal diet and sex of animals (p < 0.0001).

**Table 1.** The impact of maternal dietary deficiencies in folic acid or choline on offspring fecal metabolite levels prior to ischemic stroke and summary of two-way ANOVA results.

| Metabolite | Sex of mice | CD | FADD | ChDD | *p*-values Interaction<br>Sex Maternal diet |
| --- | --- | --- | --- | --- | --- |
| HEPES | Females | 5.56 ± 0.04 | 5.29 ± 0.02 | 6.41 ± 0.22 | 0.01 |
|  | Males | 5.68 ± 0.20 | 5.39 ± 0.04 | 5.41 ± 0.03 | 0.994 |
|  |  |  |  |  | 0.00052 |

Values are expressed as the mean ± SEM of 5 to 7 mice per group. Abbreviations: CD: Control diet; ChDD: choline-deficient diet; FADD: folic acid-deficient diet.

### One-Week Post-Stroke

#### Interactions between maternal diet and sex

There were interactions between maternal diet and offspring sex in PALDA (Table 2; p = 0.213), 3-Oxopalmitic Acid (p = 0.001), 7-Dehydrocholesterol (p = 0.001), Tetrahydrodeoxycorticosterone (p = 0.001), and Avenasterol (p = 0.011).

**Table 2.**
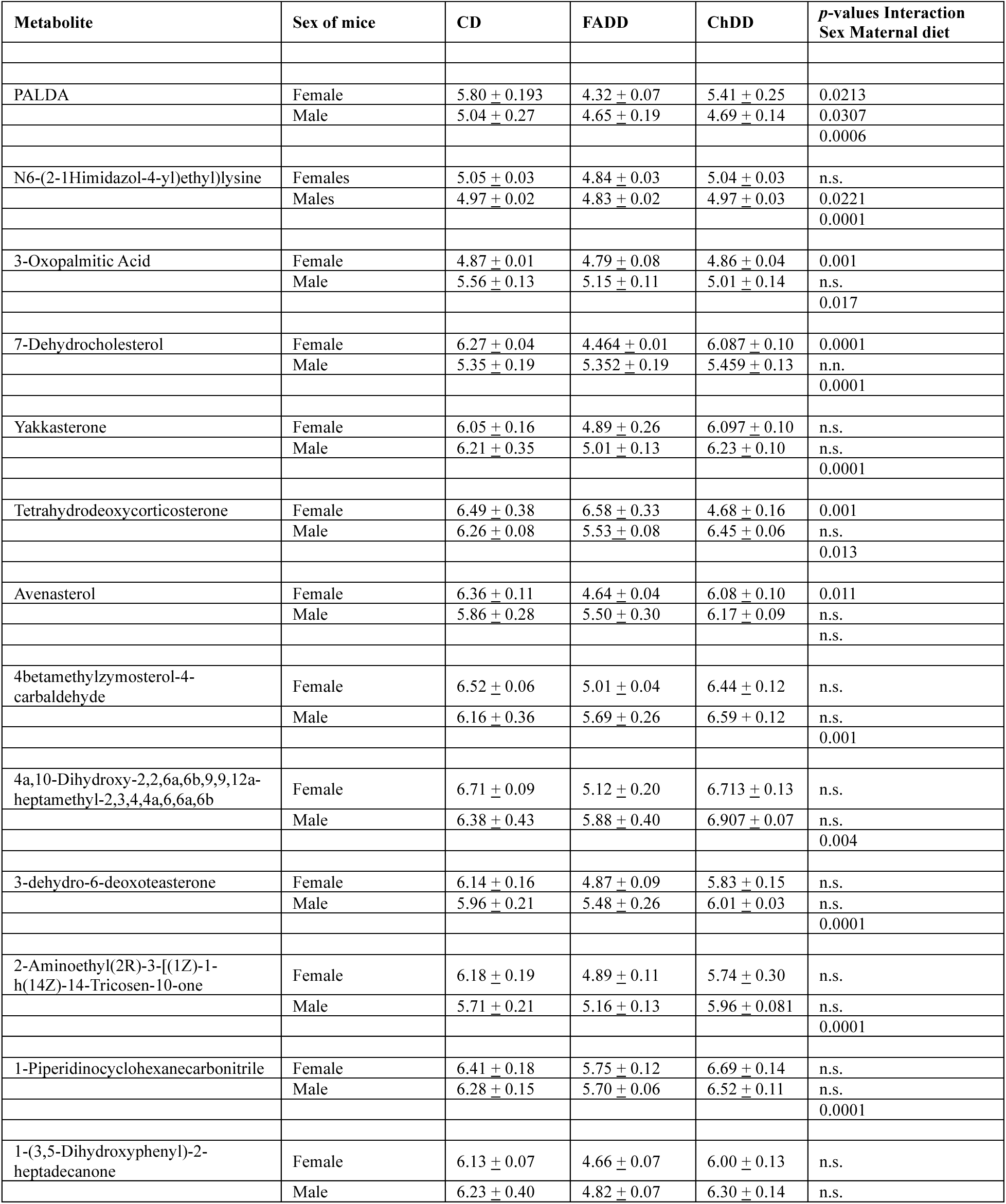
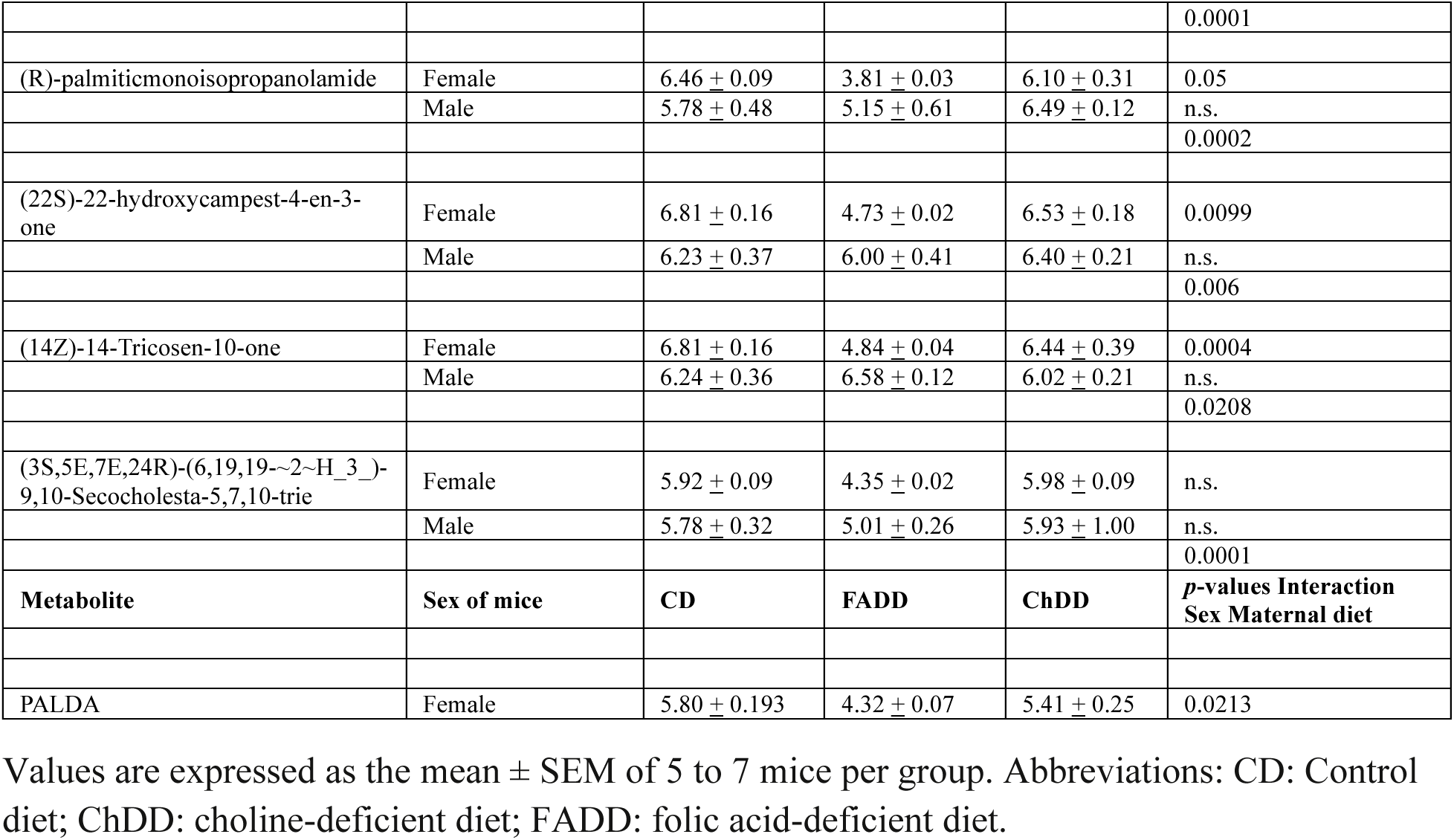
The impact of maternal dietary deficiencies in folic acid or choline on offspring fecal metabolite levels 1 week post ischemic stroke and summary of two-way ANOVA results.

| Metabolite | Sex of mice | CD | FADD | ChDD | p-values Interaction<br>Sex Maternal diet |
| --- | --- | --- | --- | --- | --- |
| PALDA | Female | 5.80 ± 0.193 | 4.32 ± 0.07 | 5.41 ± 0.25 | 0.0213 |
|  | Male | 5.04 ± 0.27 | 4.65 ± 0.19 | 4.69 ± 0.14 | 0.0307 |
|  |  |  |  |  | 0.0006 |
| N6-(2-Himidazol-4-yl)ethyl)lysine | Females | 5.05 ± 0.03 | 4.84 ± 0.03 | 5.04 ± 0.03 | n.s. |
|  | Males | 4.97 ± 0.02 | 4.83 ± 0.02 | 4.97 ± 0.03 | 0.0221 |
|  |  |  |  |  | 0.0001 |
| 3-Oxopalmitic Acid | Female | 4.87 ± 0.01 | 4.79 ± 0.08 | 4.86 ± 0.04 | 0.001 |
|  | Male | 5.56 ± 0.13 | 5.15 ± 0.11 | 5.01 ± 0.14 | n.s. |
|  |  |  |  |  | 0.017 |
| 7-Dehydrocholesterol | Female | 6.27 ± 0.04 | 4.464 ± 0.01 | 6.087 ± 0.10 | 0.0001 |
|  | Male | 5.35 ± 0.19 | 5.352 ± 0.19 | 5.459 ± 0.13 | n.n. |
|  |  |  |  |  | 0.0001 |
| Yakkasterone | Female | 6.05 ± 0.16 | 4.89 ± 0.26 | 6.097 ± 0.10 | n.s. |
|  | Male | 6.21 ± 0.35 | 5.01 ± 0.13 | 6.23 ± 0.10 | n.s. |
|  |  |  |  |  | 0.0001 |
| Tetrahydrodeoxycorticosterone | Female | 6.49 ± 0.38 | 6.58 ± 0.33 | 4.68 ± 0.16 | 0.001 |
|  | Male | 6.26 ± 0.08 | 5.53 ± 0.08 | 6.45 ± 0.06 | n.s. |
|  |  |  |  |  | 0.013 |
| Avenasterol | Female | 6.36 ± 0.11 | 4.64 ± 0.04 | 6.08 ± 0.10 | 0.011 |
|  | Male | 5.86 ± 0.28 | 5.50 ± 0.30 | 6.17 ± 0.09 | n.s. |
|  |  |  |  |  | n.s. |
| 4betamethylzymosterol-4-carbaldehyde | Female | 6.52 ± 0.06 | 5.01 ± 0.04 | 6.44 ± 0.12 | n.s. |
|  | Male | 6.16 ± 0.36 | 5.69 ± 0.26 | 6.59 ± 0.12 | n.s. |
|  |  |  |  |  | 0.001 |
| 4a,10-Dihydroxy-2,2,6a,6b,9,9,12a-heptamethyl-2,3,4,4a,6,6a,6b | Female | 6.71 ± 0.09 | 5.12 ± 0.20 | 6.713 ± 0.13 | n.s. |
|  | Male | 6.38 ± 0.43 | 5.88 ± 0.40 | 6.907 ± 0.07 | n.s. |
|  |  |  |  |  | 0.004 |
| 3-dehydro-6-deoxoteasterone | Female | 6.14 ± 0.16 | 4.87 ± 0.09 | 5.83 ± 0.15 | n.s. |
|  | Male | 5.96 ± 0.21 | 5.48 ± 0.26 | 6.01 ± 0.03 | n.s. |
|  |  |  |  |  | 0.0001 |
| 2-Aminoethyl(2R)-3-[(1Z)-1-h(14Z)-14-Tricosen-10-one | Female | 6.18 ± 0.19 | 4.89 ± 0.11 | 5.74 ± 0.30 | n.s. |
|  | Male | 5.71 ± 0.21 | 5.16 ± 0.13 | 5.96 ± 0.081 | n.s. |
|  |  |  |  |  | 0.0001 |
| 1-Piperidinocyclohexanecarbonitrile | Female | 6.41 ± 0.18 | 5.75 ± 0.12 | 6.69 ± 0.14 | n.s. |
|  | Male | 6.28 ± 0.15 | 5.70 ± 0.06 | 6.52 ± 0.11 | n.s. |
|  |  |  |  |  | 0.0001 |
| 1-(3,5-Dihydroxyphenyl)-2-heptadecanone | Female | 6.13 ± 0.07 | 4.66 ± 0.07 | 6.00 ± 0.13 | n.s. |
|  | Male | 6.23 ± 0.40 | 4.82 ± 0.07 | 6.30 ± 0.14 | n.s. |
|  |  |  |  |  | 0.0001 |
| (R)-palmiticmonoisoopropanolamide | Female | 6.46 ± 0.09 | 3.81 ± 0.03 | 6.10 ± 0.31 | 0.05 |
|  | Male | 5.78 ± 0.48 | 5.15 ± 0.61 | 6.49 ± 0.12 | n.s. |
|  |  |  |  |  | 0.0002 |
| (22S)-22-hydroxycampest-4-en-3-one | Female | 6.81 ± 0.16 | 4.73 ± 0.02 | 6.53 ± 0.18 | 0.0099 |
|  | Male | 6.23 ± 0.37 | 6.00 ± 0.41 | 6.40 ± 0.21 | n.s. |
|  |  |  |  |  | 0.006 |
| (14Z)-14-Tricosen-10-one | Female | 6.81 ± 0.16 | 4.84 ± 0.04 | 6.44 ± 0.39 | 0.0004 |
|  | Male | 6.24 ± 0.36 | 6.58 ± 0.12 | 6.02 ± 0.21 | n.s. |
|  |  |  |  |  | 0.0208 |
| (3S,5E,7E,24R)-(6,19,19~2~H_3)-9,10-Secocholesta-5,7,10-trie | Female | 5.92 ± 0.09 | 4.35 ± 0.02 | 5.98 ± 0.09 | n.s. |
|  | Male | 5.78 ± 0.32 | 5.01 ± 0.26 | 5.93 ± 1.00 | n.s. |
|  |  |  |  |  | 0.0001 |
| **Metabolite** | **Sex of mice** | **CD** | **FADD** | **ChDD** | **p-values Interaction**<br>**Sex Maternal diet** |
| PALDA | Female | 5.80 ± 0.193 | 4.32 ± 0.07 | 5.41 ± 0.25 | 0.0213 |

Values are expressed as the mean ± SEM of 5 to 7 mice per group. Abbreviations: CD: Control diet; ChDD: choline-deficient diet; FADD: folic acid-deficient diet.

### Sex differences

Sex differences were observed in in PALDA (Table 2, p = 0.0307) and N6-(2-1Himidazol- 4-yl) ethyl) lysine (p = 0.0221). Significant sex differences were followed up with pairwise comparisons, details are listed in Table 2.

### Impact of Maternal Diet

After one-week post-stroke we observed maternal dietary differences in fecal measurements for the following metabolites, PALDA (Table 2; p = 0.0006), N6-(2-1Himidazol- 4-yl) ethyl)lysine (p = 0.001), 3-Oxopalmitic Acid (p = 0.017), 7-Dehydrocholesterol (p = 0.0001), Yakkasterone (p = 0.0001), Tetrahydrodeoxycorticosterone (p = 0.013), Cromide BEM (p=0.0063), certonardosterol (p = 0.0001), 4betamethylzymosterol-4-carbaldehyde (p = 0.001), 4a,10-Dihydroxy-2,2,6a,6b,9,9,12a-heptamethyl-2,3,4,4a,6,6a,6b (p = 0.04), 3-dehydro-6- deoxoteasterone (p = 0.001), 2-Aminoethyl(2R)-3-[(1Z)-1-h(14Z)-14-Tricosen-10-one (p = 0.0001), 2-Aminoethyl(2R)-3-[(1Z)-1-h(14Z)-14-Tricosen-10-one (p = 0.0001), 1-(3,5-Dihydroxyphenyl)-2-heptadecanone (p = 0.0001), (R)-palmiticmonoisopropanolamide (p = 0.0002), (R)-palmiticmonoisopropanolamide (p = 0.006), and (3S,5E,7E,24R)-(6,19,19-∼2∼H_3_)-9,10-Secocholesta-5,7,10-trie (p = 0.0001). Significant maternal diet differences in metabolites were followed up with pairwise comparisons, details are listed in table 2.

### Four-Week Post-Stroke

#### Interactions between maternal diet and sex

There was an interaction between maternal diet and offspring sex for the following metabolites, HEPES (Table 3, p = 0.032), 1-(4-Aminobutyl)urea (p = 0.0015), PRIMA-1 (p = 0.0005), N-lauroylglycine (p = 0.0036), N∼6∼,N∼6∼-Dimethyllysine (p = 0.0088), Misoprostol (p = 0.032), Gly-Leu (p = 0.003), Chamazulene (p = 0.02), Esmolol (p = 0.013), Dodecylethanolamide (p = 0.038), Creatine (p = 0.01), C2 Dihydroceramide (p = 0.003), BAR501 (p = 0.006), APM (p = 0.024), Adenine (p = 0.012), Acetylcadaverine (p = 0.0038), Acetohydroxamic acid (p = 0.0084), 1-PP (p = 0.011), (4R)-5-Hydroxy-L-leucine (p = 0.003), and (+)-castanospermine (p = 0.028).

**Table 3.** The impact of maternal dietary deficiencies in folic acid or choline on offspring fecal metabolite levels 4 weeks post ischemic stroke and summary of two-way ANOVA results.

| Metabolite | Sex of mice | CD | FADD | ChDD | *p*-values Interaction Sex Maternal diet |
| --- | --- | --- | --- | --- | --- |
| HEPES | Females | 5.27 ± 0.03 | 5.41 ± 0.02 | 5.50 ± 0.06 | 0.032 |
|  | Males | 5.39 ± 0.03 | 5.44 ± 0.04 | 5.42 ± 0.03 | n.s. |
|  |  |  |  |  | 0.0072 |
| Fenamole | Female | 5.89 ± 0.14 | 6.52 ± 0.067 | 6.12 ± 0.085 | n.s. |
|  | Male | 6.46 ± 0.12 | 5.92 ± 0.058 | 6.37 ± 0.042 | 0.03 |
|  |  |  |  |  | 0.02 |
| 2,3-Dihydroxypropylstearate | Female | 6.43 ± 0.21 | 6.41 ± 0.12 | 6.32 ± 0.21 | n.s. |
|  | Male | 6.24 ± 0.15 | 6.13 ± 0.07 | 6.27 ± 0.11 | n.s. |
|  |  |  |  |  | 0.0004 |
| n-Hexanamide | Female | 5.89 ± 0.29 | 6.69 ± 0.04 | 6.67 ± 0.02 | 0.0073 |
|  | Male | 7.25 ± 0.13 | 7.25 ± 0.35 | 6.57 ± 0.30 | 0.0019 |
|  |  |  |  |  | n.s. |
| N-Acetyl-Lglutamic Acid | Female | 7.01 ± 0.15 | 7.27 ± 0.12 | 7.24 ± 0.10 | 0.0029 |
|  | Male | 7.75 ± 0.11 | 7.18 ± 0.11 | 7.20 ± 0.13 | 0.051 |
|  |  |  |  |  | n.s. |
| JWH 133 | Female | 6.48 ± 0.29 | 6.68 ± 0.08 | 6.75 ± 0.07 | n.s. |
|  | Male | 6.63 ± 0.06 | 6.68 ± 0.07 | 6.74 ± 0.05 | n.s. |
|  |  |  |  |  | <0.0001 |
| Homocitrulline | Female | 5.68 ± 0.03 | 6.27 ± 0.09 | 5.91 ± 0.07 | 0.0001 |
|  | Male | 6.32 ± 0.08 | 5.72 ± 0.12 | 5.94 ± 0.16 | n.s. |
|  |  |  |  |  | n.s. |
| Sphinganine | Female | 6.73 ± 0.17 | 7.14 ± 0.03 | 7.07 ± 0.03 | 0.018 |
|  | Male | 7.16 ± 0.03 | 7.11 ± 0.03 | 7.07 ± 0.01 | 0.02 |
|  |  |  |  |  | 0.053 |
| R-(+)-Etiracetam | Female | $5.21 \pm 0.22$ | $5.99 \pm 0.04$ | $5.71 \pm 0.05$ | 0.0005 |
| | Male | $5.81 \pm 0.06$ | $5.60 \pm 0.10$ | $5.57 \pm 0.09$ | n.s. |
|  |  |  |  |  | n.s. |
| Pyrrolidine | Female | $5.59 \pm 0.10$ | $6.08 \pm 0.12$ | $5.75 \pm 0.12$ | 0.01 |
| | Male | $6.14 \pm 0.09$ | $5.91 \pm 0.12$ | $5.93 \pm 0.10$ | 0.04 |
|  |  |  |  |  | n.s. |
| 1-(4-Aminobutyl)urea | Female | $4.61 \pm 0.19$ | $5.94 \pm 0.08$ | $5.44 \pm 0.17$ | 0.0015 |
| | Male | $5.40 \pm 0.09$ | $5.15 \pm 0.34$ | $5.30 \pm 0.24$ | n.s. |
|  |  |  |  |  | 0.025 |
| Propoxur | Female | $5.31 \pm 0.19$ | $6.017 \pm 0.07$ | $5.93 \pm 0.07$ | n.s. |
| | Male | $5.94 \pm 0.14$ | $6.072 \pm 0.09$ | $6.18 \pm 0.14$ | 0.006 |
|  |  |  |  |  | 0.003 |
| PRIMA-1 | Female | $5.20 \pm 0.22$ | $6.00 \pm 0.04$ | $5.71 \pm 0.05$ | 0.0005 |
| | Male | $5.81 \pm 0.06$ | $5.60 \pm 0.10$ | $5.57 \pm 0.09$ | n.s. |
|  |  |  |  |  | n.s. |
| Phytosphingosine | Female | $7.56 \pm 0.21$ | $8.14 \pm 0.19$ | $8.02 \pm 0.07$ | 0.022 |
| | Male | $8.13 \pm 0.07$ | $8.13 \pm 0.018$ | $8.15 \pm 0.02$ | 0.012 |
|  |  |  |  |  | 0.02 |
| N-lauroylglycine | Female | $5.96 \pm 0.13$ | $6.46 \pm 0.04$ | $6.50 \pm 0.09$ | 0.0036 |
| | Male | $6.34 \pm 0.09$ | $6.22 \pm 0.11$ | $6.32 \pm 0.04$ | n.s. |
|  |  |  |  |  | 0.021 |
| Nicaraven | Female | $5.24 \pm 0.09$ | $4.88 \pm 0.07$ | $4.84 \pm 0.05$ | n.s. |
| | Male | $5.06 \pm 0.07$ | $5.02 \pm 0.10$ | $4.94 \pm 0.03$ | n.s. |
|  |  |  |  |  | 0.003 |
| N~6~,N~6~-<br>Dimethyllysine | Female | $5.05 \pm 0.14$ | $5.78 \pm 0.07$ | $5.47 \pm 0.13$ | 0.0088 |
| | Male | $5.45 \pm 0.12$ | $5.31 \pm 0.16$ | $5.63 \pm 0.18$ | n.s. |
|  |  |  |  |  | 0.049 |
| Misoprostol | Female | $6.18 \pm 0.14$ | $5.85 \pm 0.08$ | $5.69 \pm 0.05$ | 0.032 |
| | Male | $5.88 \pm 0.05$ | $5.82 \pm 0.11$ | $5.91 \pm 0.10$ | n.s. |
|  |  |  |  |  | 0.036 |
| Lovastatin | Female | $6.86 \pm 0.13$ | $6.43 \pm 0.10$ | $6.52 \pm 0.05$ | n.s. |
| | Male | $6.61 \pm 0.04$ | $6.50 \pm 0.05$ | $6.53 \pm 0.08$ | n.s. |
|  |  |  |  |  | 0.0081 |
| HMMNI | Female | $4.87 \pm 0.19$ | $5.48 \pm 0.23$ | $5.50 \pm 0.05$ | n.s. |
| | Male | $6.08 \pm 0.19$ | $6.03 \pm 0.19$ | $5.78 \pm 0.44$ | 0.0014 |
|  |  |  |  |  | n.s. |
| Gly-Leu | Female | $5.38 \pm 0.10$ | $6.46 \pm 0.13$ | $5.96 \pm 0.11$ | 0.003 |
| | Male | $6.21 \pm 0.15$ | $6.07 \pm 0.30$ | $5.88 \pm 0.20$ | n.s. |
|  |  |  |  |  | 0.026 |
| Chamazulene | Female | $5.78 \pm 0.35$ | $5.01 \pm 0.01$ | $4.929 \pm 0.02$ | 0.02 |
| | Male | $4.99 \pm 0.03$ | $4.97 \pm 0.03$ | $4.99 \pm 0.02$ | 0.05 |
|  |  |  |  |  | 0.02 |
| Esmolol | Female | $6.00 \pm 0.22$ | $6.39 \pm 0.07$ | $6.43 \pm 0.04$ | 0.013 |
| | Male | $6.59 \pm 0.03$ | $6.51 \pm 0.08$ | $6.35 \pm 0.06$ | 0.025 |
|  |  |  |  |  | 0.0364 |
| Dodecylethanolamide | Female | $6.38 \pm 0.09$ | $6.15 \pm 0.03$ | $6.16 \pm 0.01$ | 0.038 |
| | Male | $6.24 \pm 0.02$ | $6.23 \pm 0.03$ | $6.20 \pm 0.02$ | n.s. |
|  |  |  |  |  | 0.009 |
| Creatine | Female | $7.21 \pm 0.17$ | $6.86 \pm 0.20$ | $7.06 \pm 0.14$ | 0.01 |
| | Male | $6.21 \pm 0.22$ | $6.98 \pm 0.10$ | $7.15 \pm 0.33$ | n.s. |
|  |  |  |  |  | n.s. |
| C2 Dihydroceramide | Female | $5.55 \pm 0.09$ | $5.87 \pm 0.06$ | $5.88 \pm 0.03$ | 0.003 |
| | Male | $5.93 \pm 0.04$ | $5.73 \pm 0.13$ | $5.84 \pm 0.05$ | n.s. |
|  |  |  |  |  | n.s. |
| BAR501 | Female | $4.61 \pm 0.06$ | $5.15 \pm 0.04$ | $4.96 \pm 0.09$ | 0.006 |
| | Male | $5.06 \pm 0.08$ | $4.99 \pm 0.11$ | $4.97 \pm 0.15$ | n.s. |
|  |  |  |  |  | 0.042 |
| APM | Female | $5.59 \pm 0.26$ | $6.42 \pm 0.07$ | $6.21 \pm 0.10$ | 0.024 |
| | Male | $6.16 \pm 0.12$ | $6.13 \pm 0.14$ | $6.14 \pm 0.13$ | n.s. |
|  |  |  |  |  | 0.038 |
| Adenine | Female | $6.51 \pm 0.14$ | $7.04 \pm 0.20$ | $6.91 \pm 0.21$ | 0.012 |
| | Male | $7.39 \pm 0.16$ | $6.76 \pm 0.19$ | $7.00 \pm 0.15$ | n.s. |
|  |  |  |  |  | n.s. |
| Acetylcadaverine | Female | $5.31 \pm 0.01$ | $5.80 \pm 0.16$ | $5.42 \pm 0.05$ | 0.0038 |
| | Male | $5.58 \pm 0.07$ | $5.44 \pm 0.06$ | $5.38 \pm 0.03$ | n.s. |
|  |  |  |  |  | 0.041 |
| Acetohydroxamic acid | Female | $7.31 \pm 0.08$ | $7.75 \pm 0.07$ | $7.48 \pm 0.09$ | 0.0084 |
| | Male | $7.83 \pm 0.12$ | $7.53 \pm 0.11$ | $7.73 \pm 0.17$ | n.s. |
|  |  |  |  |  | n.s. |
| 5-Hydroxytryptophol | Female | $5.32 \pm 0.14$ | $6.02 \pm 0.07$ | $5.93 \pm 0.07$ | n.s. |
| | Male | $5.94 \pm 0.19$ | $6.07 \pm 0.09$ | $6.18 \pm 0.14$ | 0.006 |
|  |  |  |  |  | 0.003 |
| 1-PP | Female | $5.61 \pm 0.08$ | $6.07 \pm 0.08$ | $5.79 \pm 0.10$ | 0.011 |
| | Male | $6.14 \pm 0.13$ | $5.85 \pm 0.12$ | $6.01 \pm 0.16$ | n.s. |
|  |  |  |  |  | n.s. |
| (4R)-5-Hydroxy-L-leucine | Female | $5.91 \pm 0.16$ | $6.66 \pm 0.10$ | $6.20 \pm 0.13$ | 0.003 |
| | Male | $6.64 \pm 0.13$ | $6.25 \pm 0.25$ | $6.37 \pm 0.08$ | n.s. |
|  |  |  |  |  | n.s. |
| (+)-castanospermine | Female | $5.05 \pm 0.29$ | $5.94 \pm 0.08$ | $5.59 \pm 0.10$ | 0.028 |
| | Male | $5.60 \pm 0.08$ | $5.59 \pm 0.10$ | $5.69 \pm 0.14$ | n.s. |
|  |  |  |  |  | 0.026 |

Values are expressed as the mean ± SEM of 5 to 7 mice per group. Abbreviations: CD: Control diet; ChDD: choline-deficient diet; FADD: folic acid-deficient diet.

### Sex Differences

There were sex differences in the following metabolites, Fenamole (Table 3, p = 0.03), n- Hexanamide (p = 0.0019), N-Acetyl-Lglutamic Acid (p = 0.051), Sphinganine (p = 0.02), Pyrrolidine (p = 0.04), Propoxur (p = 0.006), Phytosphingosine (p = 0.012), HMMNI (p = 0.0014), Chamazulene (p = 0.05), Esmolol (p = 0.025), and 5-Hydroxytryptophol (p = 0.006). There were interactions between maternal diet and sex in the following metabolites, n-Hexanamide (p = 0.0073), N-Acetyl-Lglutamic Acid (p = 0.0029), Homocitrulline (p = 0.0001), Sphinganine (p = 0.018), R-(+)-Etiracetam (p = 0.0005), and Pyrrolidine (p = 0.01). Significant sex differences between females and males were followed up with pair wise comparisons, details are listed in table 4.

**Table 4.** One-week post-stroke Tukey’s pairwise comparison summary determining the impact of maternal dietary deficiencies in folic acid or choline on offspring fecal metabolite levels.

| Metabolite | Female |  |  | Male |  |  |
| --- | --- | --- | --- | --- | --- | --- |
|  | CD vs. FADD | CD vs. ChDD | ChDD vs. FADD | CD vs. FADD | CD vs. ChDD | ChDD vs. FADD |
| PALDA | 0.002 | n.s. | 0.0045 | n.s. | n.s. | n.s. |
| N6-(2-Imidazol-4-yl)ethyllysine | 0.0002 | n.s. | 0.0003 | 0.0044 | n.s. | 0.0002 |
| 3-Oxopalmitic Acid | n.s. | n.s. | n.s. | < 0.001 | < 0.001 | n.s. |
| 7-Dehydrocholesterol | < 0.001 | < 0.001 | n.s. | n.s. | n.s. | n.s. |
| Yakkasterone | 0.0013 | n.s. | 0.001 | 0.003 | n.s. | 0.0001 |
| Tetrahydrodeoxycorticosterone | n.s. | 0.0002 | 0.0001 | n.s. | n.s. | 0.0276 |
| Avenasterol | 0.001 | n.s. | 0.0003 | n.s. | n.s. | n.s. |
| 4betamethylzymosterol-4-carbaldehyde | 0.0004 | n.s. | 0.0007 | n.s. | n.s. | 0.0004 |
| 4a,10-Dihydroxy-2,2,6a,6b,9,9,12a-heptamethyl-2,3,4,4a,6,6a,6b | 0.0034 | n.s. | 0.003 | n.s. | n.s. | 0.0354 |
| 3-dehydro-6-deoxoteasterone | 0.0003 | n.s. | 0.0043 | n.s. | n.s. | n.s. |
| 2-Aminoethyl(2R)-3-[(1Z)-1-h(14Z)-14-Tricosen-10-one | 0.0008 | n.s. | 0.0016 | n.s. | n.s. | 0.0306 |
| 1-Piperidinocyclohexanecarbonitrile | 0.0085 | n.s. | 0.0002 | 0.0003 | n.s. | 0 |
| 1-(3,5-Dihydroxyphenyl)-2-heptadecanone | 0.003 | n.s. | 0.0001 | 0.0009 | n.s. | n.s. |
| (R)-palmiticmonoisopropanolamide | 0.004 | n.s. | 0.0004 | n.s. | n.s. | 0.0017 |
| (22S)-22-hydroxycampest-4-en-3-one | 0.0002 | n.s. | 0.0006 | n.s. | n.s. | n.s. |
| (14Z)-14-Tricosen-10-one | 0.0002 | n.s. | 0.0018 | n.s. | n.s. | n.s. |
| (3S,5E,7E,24R)-(6,19,19~2~H_3)-9,10-Secocholesta-5,7,10-trie | 0.0255 | n.s. | 0.007 | 0.0001 | n.s. | < 0.001 |

Abbreviations: CD: Control diet; ChDD: choline-deficient diet; FADD: folic acid-deficient diet.

### Impact of Maternal Diet

At four weeks post-stroke there were differences in offspring metabolites, as a result of maternal diet in the following: HEPES (Table 3, p = 0.0072), Fenamole (p = 0.03), 2,3-Dihydroxypropylstearate (p = 0.0004), JWH 133 (p < 0.0001), Sphinganine (p = 0.053), 1-(4- Aminobutyl)urea (p = 0.025), Propoxur (p = 0.003), phytosphingosine (p = 0.02), N-lauroylglycine (p = 0.021), nicaraven (p = 0.003), N∼6∼,N∼6∼-Dimethyllysine (p = 0.049), Misoprostol (p = 0.036), lovastatin (p = 0.0081), Gly-Leu (p=0.026), Chamazulene (p = 0.02), Esmolol (p = 0.0364), Dodecylethanolamide (p = 0.009), BAR501(p = 0.042), APM (p = 0.038), 5-Hydroxytryptophol (p = 0.003), and (+)-castanospermine (p = 0.026). Significant differences in metabolite levels as a result of maternal diet was followed up with pairwise comparisons, details are listed in table 5.

**Table 5.**
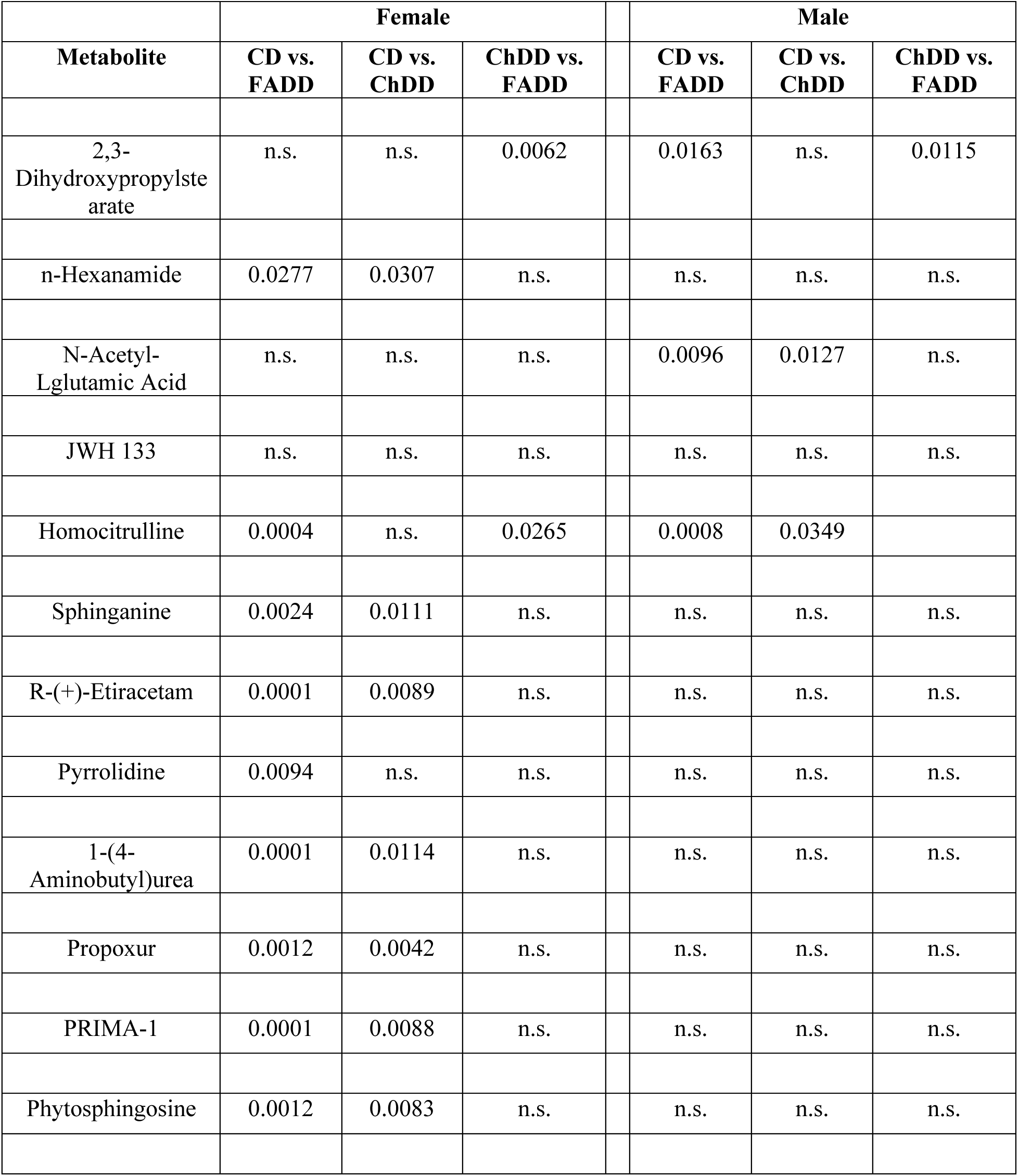
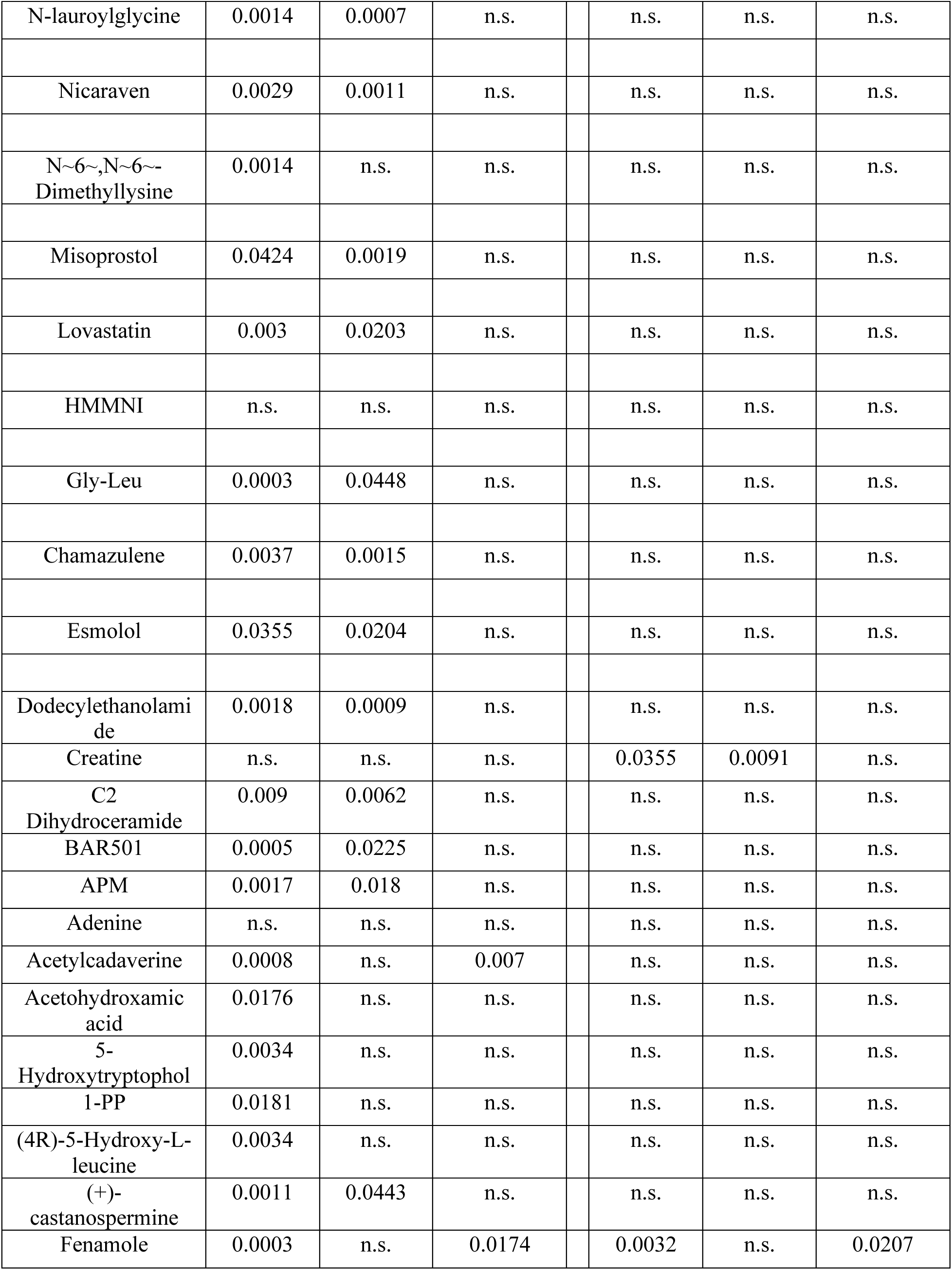
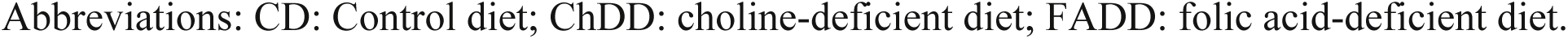
Four-week post-stroke Tukey’s pairwise comparison summary determining the impact of maternal dietary deficiencies in folic acid or choline on offspring fecal metabolite levels.

## Discussion

Ischemic stroke is a major health concern, with nutrition being a modifiable risk factor. We have previously demonstrated offspring from female mice deficient in either folic acid or choline have worse outcome after stroke (Clementson et al., 2022; Hurley et al., 2023; Pull et al., 2023) . Metabolites are important marker to measure the changes in the gut microbiota, which can be significant markers for different pathophysiological diseases in the body. We collected fecal samples from female and male offspring whose mothers were maintained on either FADD, ChDD, and CD. The fecal samples were collected at three timepoints including prior to, 1, and 4 weeks post-ischemic stroke to measure metabolites using untargeted analyses. This untargeted metabolomics analysis revealed a wide spectrum of detrimental effects caused by stroke and maternal diet deficiencies in 3-month-old offspring. Our analysis revealed that maternal folic acid dietary deficiency has significant impact on microbiota of offspring after ischemic stroke. The results demonstrate that sex and maternal diet impact metabolites levels after ischemic stroke and may impact outcomes.

At pre-stroke stage we observed changes in levels of HEPES because of maternal diet. The role and involvement of these metabolites remain unclear. The differences in maternal diet and sex impact on the stroke outcome become more prominent in 1-and 4-week post-stroke time points.

A total of 17 metabolites differences observed in different diet groups at 1 week post stroke. FADD mice had lower level of metabolites compared to ChDD and CD, while ChDD mice had similar metabolites level to CD mice. Male mice had lower levels of metabolites at 1 week post stroke. Most of the observed metabolites observed in lower levels are involved in various survival and developmental cellular mechanisms including cholesterol biogenesis, cell division, macrophage polarization, and neuroprotection. Tertahydrodeoxycorticsterone, a neurosteroid, have been observed to play a role in neuroprotection and neural analgesic affects (Kang et al., 2021; Márk & Paragh, 2007; Mediratta et al., 2001; Patchev et al., 1997; Piette, 2020; Rupprecht, 1997; Wetzel et al., 1999; Womack et al., 2006). Another metabolite, 7-dehydrocholesterol reductase (DHCR7) is an enzyme that catalyzes 7-dehydrocholesterol (7-DHC) to cholesterol, a final step in the biogenesis of cholesterol (Huang et al., 2017; Waterham & Wanders, 2000; Xiao et al., 2020). DHCR7 mutations in humans cause a clinical autosomal recessive genetic disease, namely Smith-Lemli-Opitz syndrome (SLOS), which is characterized with multiple abnormalities including growth deficiency, intellectual disability, and frequent infections(Honda et al., 1998). JWH 133 activates cannabinoid 2 receptors in reduced ovarian ischemia-reperfusion injury due to its antioxidant and anti-inflammatory effects (Aydogan Kirmizi et al., 2021; López-Dyck et al., 2017; Onat et al., 2021; Wojcieszak et al., 2016). JWH-133 showed anti-obesity effects that ameliorated pro-inflammatory M1 macrophage polarization through the Nrf2/HO-1 pathway (Q. Wu et al., 2020). The lower levels of the involved metabolites seem to have effects on neuroprotective mechanism during neurodevelopment in early age as well as have effects on the immune systems and fatty acid mechanism. Dysfunction in such metabolites can also signal towards growth deficiency and more prone to infections. However, the mechanism of the effects remains unclear and require additional investigation.

At 4-week post stroke, metabolites significantly lower compared to one-week timepoint are observed to have come back to normal levels while new metabolites were found to be in abnormal levels. Female mice were affected more adversely with lower levels of all the metabolites found in 4-week post stroke compared to pre-stroke levels. Mice on either ChDD or FADD were both at lower levels as compared mice of CD. Homocitrulline levels were lower in both genders on FADD diet which plays role in cell viability and mitochondrial function in menadione-treated astrocytes (Zanatta et al., 2016). N-Acetyl-glutamic acid, Noradrenaline, and Creatine which play roles in liver ureagenesis, cognitive process, and cellular metabolism essential for cognitive abilities respectively were found in lover levels (Caldovic et al., 2010; Harper et al., 2009; Prokopová, 2010; Sumien et al., 2018). Other metabolites, Dehydrocostus which have antibacterial activity by inhibiting LPS-induced production of proinflammatory mediators were in lower levels (Nie et al., 2019). Metabolite which are indicators of central nervous system health and play protective roles were detected at lower levels in female with both FADD and CHDD diet. Some of the metabolites are (+)-Castanospermine involved in reduces inflammation in brain, 5- Hydroxytryptophol responsible for elicits cerebro arterial contractions (Saul et al., 1983; Tharappel et al., 2020), Acetylcadaverine lower levels is markedly found Alzheimer’s Disease (Paik et al., 2006), BAR501 responsible for reversing intestinal inflammation and shifts activation of intestinal macrophages reducing expression of inflammatory genes (Biagioli et al., 2017). C2 Dihydroceramide associated with increased apoptosis in human leukemia HL-60 cells (Shikata et al., 2003), and Gly-Leu, Glycine acts as a neurotransmitter in central nervous system; antioxidant, anti-inflammatory, cryoprotective, and immunomodulatory in peripheral and nervous tissues and leucine essential amino acid used in ATP generation, protein synthesis, tissue regeneration, and metabolism (Pedroso et al., 2015; Razak et al., 2017).

Overall, after 4-week post stroke, it is strongly evident that female offspring that come from mothers maintained on FADD or ChDD have more changes in their metabolites compared to males. The changes are magnified at the four-week timepoint despite being on CD compared to 1 week post stroke. The abnormal metabolites levels suggest that there is significant inflammation found both in brain and gut. It is indicated that the body’s anti-inflammatory and defense system is also highly affected. Interestingly, markers for Alzheimer’s disease, cancers, gut, and brain inflammation were found to be changed suggesting that these subjects might experience an early physical deterioration and hindered development. In a previous study, there was evidence of phenotypical behavioral issues in mice despite any changes in the ischemic size of the brain infarct. This is suggestive of activation of multiple pathophysiological mechanisms that have long lasting effects on a developing body. This is the first study to report changes in fecal sample metabolites after changes in maternal diet and ischemic stroke. More work is required, as the mechanism and molecular pathway are not clear. Possible next steps may include sequencing samples to determine changes in DNA. The present study provides insights that maternal dietary deficiencies in folic acid when combined with ischemic stroke impact offspring metabolites in a sex dependent manner.

## Materials and methods

### Animals and experimental design

All experiments were conducted in accordance with the guidelines of the Midwestern University Institutional Animal Care Users Committee (IACUC 2983). C57/BL6J mice (both female and male) were obtained from Jackson Laboratories. The breeding pairs produced male (n = 17) and female (n = 15) offspring.

Experimental manipulations are summarized in Figure 2. Two-month-old female mice were habituated for seven days before they were placed on either control (CD), folic acid (FADD, folic acid: 0.3 mg/kg), or choline deficient diets (ChDD, choline bitrate: 300 mg/kg) (Clementson et al., 2023; Dam et al., 2017; Hurley et al., 2023; Jadavji, Deng, et al., 2015; Jadavji, Farr, et al., 2015) (Envigo, Indianapolis, IN). The control diet contains the minimum amount of folic acid (2 mg/kg) and choline bitrate (1150 mg/kg). The dams were maintained on the diets 4 weeks prior to pregnancy, during pregnancy, and lactation. Once female and male offspring were weaned from their mothers, they were maintained on a CD. At 2 months of age, the offspring were subjected to ischemic stroke, using the photothrombosis model to the sensorimotor cortex. Stool samples were collected from the same offspring at three timepoints: prior to ischemic stroke, 1 week after, and 4 weeks after the stroke.

**Figure 2.**
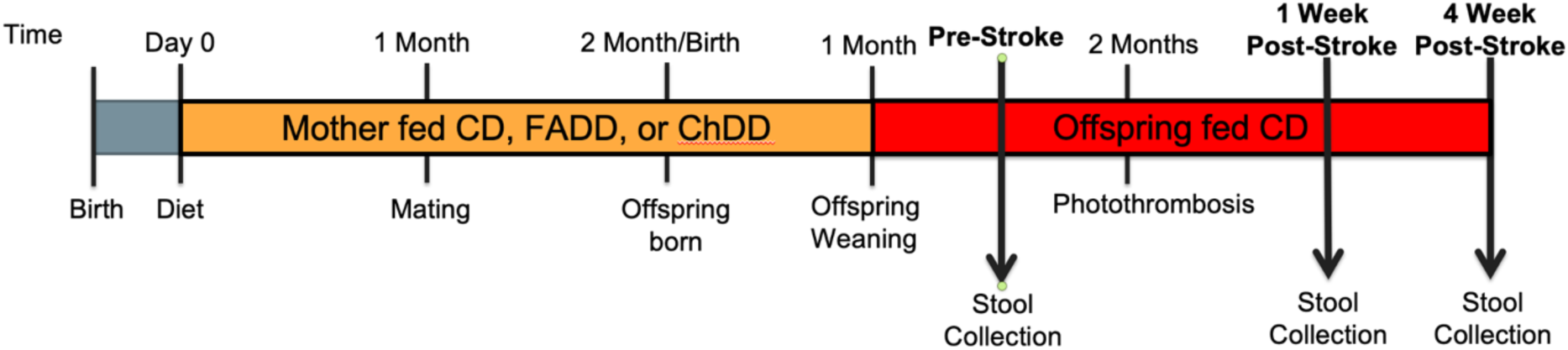
Displays the timeline of the experiment. Pregnant mothers were fed either the CD, FADD, or ChDD diet throughout the months of pregnancy and lactation until the offspring were weaned. Once the offspring were weaned, they were maintained on the CD. At 2 months of age, the offspring were subjected to ischemic stroke via the PT model. Tissue and fecal matter was collected at 1-Week Post Stroke and 4-Week Post Stroke.

### Photothrombosis Model for Ischemic Stroke

At 2 months of age all female and male offspring mice were subjected to photothrombosis to induce a unilateral ischemic stroke in the sensorimotor cortex. Mice were anesthetized with isoflurane (1.5%) in a 70:30 nitrous oxide: oxygen mixture. Core body temperature was monitored with a rectal thermometer (Harvard Apparatus, Holliston, MA) and maintained at 37 ± 0.2°C using a heating blanket. 10mg/kg of the photosensitive Rose Bengal (Sigma, St. Louis, MA) dye was injected intraperitoneally 5 minutes prior to irradiation. A 532 nm green laser was placed 3 cm above the animal and directed to the sensorimotor cortex (mediolateral + 0.24mm)(Abato et al., 2020; Clementson et al., 2023; Hurley et al., 2023; Jadavji et al., 2017, 2019, 2019; Lee et al., 2004; Mbs et al., 2023; Poole et al., 2022) for 15 min.

### Fecal preparation

Fecal samples were collected in a microfuge tube at three timepoints, prior to, 1 and 4 weeks after ischemic stroke. The samples were stored at -80°C until analysis.

Each fecal sample (∼20 mg) was homogenized in 200 µL MeOH:PBS (4:1, v:v, containing 1,810.5 μM ^13^C_3_-lactate and 142 μM ^13^C_5_-glutamic Acid) in an Eppendorf tube using a Bullet Blender homogenizer (Next Advance, Averill Park, NY). Then 800 µL MeOH:PBS (4:1, v:v, containing 1,810.5 μM ^13^C_3_-lactate and 142 μM ^13^C_5_-glutamic Acid) was added, and after vortexing for 10 s, the samples were stored at -20°C for 30 min. The samples were then sonicated in an ice bath for 30 min. The samples were centrifuged at 14,000 RPM for 10 min (4°C), and 800 µL supernatant was transferred to a new Eppendorf tube. The samples were then dried under vacuum using a CentriVap Concentrator (Labconco, Fort Scott, KS). Prior to MS analysis, the obtained residue was reconstituted in 150 μL 40% PBS/60% ACN. A quality control (QC) sample was pooled from all the study samples.

### Solutions for Experiments

Solutions used to complete analysis included, acetonitrile (ACN), methanol (MeOH), ammonium acetate, and acetic acid, all LC-MS grade, were purchased from Fisher Scientific (Pittsburgh, PA). Ammonium hydroxide was bought from Sigma-Aldrich (Saint Louis, MO). DI water was provided in-house by a Water Purification System from EMD Millipore (Billerica, MA). PBS was bought from GE Healthcare Life Sciences (Logan, UT). The standard compounds corresponding to the measured metabolites were purchased from Sigma-Aldrich (Saint Louis, MO) and Fisher Scientific (Pittsburgh, PA).

### Untargeted LC-MS Metabolomics

The untargeted LC-MS metabolomics method used has developed and used in a growing number of studies (Garcia et al., 2024; Gu et al., 2015; D. Scieszka et al., 2023; D. P. Scieszka et al., 2023; Wei et al., 2021). Briefly, all LC-MS experiments were performed using the Thermo Vanquish UPLC-Exploris 240 Orbitrap MS instrument (Waltham, MA). Fecal samples were injected twice, 10 µL for analysis using negative ionization mode and 4µL for analysis using positive ionization mode. Both chromatographic separations were performed in hydrophilic interaction chromatography (HILIC) mode on a Waters XBridge BEH Amide column (150 x 2.1 mm, 2.5 µm particle size, Waters Corporation, Milford, MA). The flow rate was 0.3 mL/min, auto- sampler temperature was kept at 4 °C, and the column compartment was set at 40 °C. The mobile phase was composed of Solvents A (10 mM ammonium acetate, 10 mM ammonium hydroxide in 95% H_2_O/5% ACN) and B (10 mM ammonium acetate, 10 mM ammonium hydroxide in 95% ACN/5% H_2_O). After the initial 1 min isocratic elution of 90% B, the percentage of Solvent B decreased to 40% at t = 11 min. The composition of Solvent B maintained at 40% for 4 min (t=15 min), and then the percentage of B gradually went back to 90%, to prepare for the next injection. Using mass spectrometer equipped with an electrospray ionization (ESI) source, we will collect untargeted data from 70 to 1050 m/z.

To identify peaks from the MS spectra, we made extensive use of the in-house chemical standards (∼600 aqueous metabolites), and in addition, we searched the resulting MS spectra against the HMDB library, Lipidmap database, METLIN database, as well as commercial databases including mzCloud, Metabolika, and ChemSpider. The absolute intensity threshold for the MS data extraction was 1,000, and the mass accuracy limit was set to 5 ppm. Identifications and annotations used available data for retention time (RT), exact mass (MS), MS/MS fragmentation pattern, and isotopic pattern. Thermo Compound Discoverer 3.3 software was used for aqueous metabolomics data processing. The untargeted data were processed by the software for peak picking, alignment, and normalization. To improve rigor, only the signals/peaks with CV < 20% across quality control (QC) pools, and the signals showing up in >80% of all the samples were included for further analysis.

### Data Preprocessing

Raw data files generated from the mass spectrometer were converted to a matrix of peak intensities, with each row representing a sample and each column representing a metabolite feature. Quality control samples were also included in the matrix for inter-batch normalization. Raw data were normalized to the median of quality control samples.

### Multivariate Analysis

Principal component analysis (PCA) and orthogonal partial least squares-discriminant analysis (OPLS-DA) were performed to explore the separation of different groups and identify important metabolites contributing to the separation.

To investigate metabolite differences between sex and maternal dietary groups, as well as their interactions, two-way ANOVA was conducted using GraphPad Prism 10. Significant main effects of two-way ANOVAs were followed by Tukey’s post-hoc test to adjust for multiple comparisons.

Statistical analysis was performed using R (version 4.1.1). For each metabolite, normality was assessed using the Shapiro-Wilk test, which is suitable for small to medium sample sizes and provides a good balance between sensitivity and specificity.

If the data for a metabolite followed a normal distribution (p > 0.05 in the Shapiro-Wilk test), independent t-tests were used to compare the means between groups. If the data did not follow a normal distribution (p ≤ 0.05 in the Shapiro-Wilk test), non-parametric Wilcoxon rank-sum tests were employed to compare the medians between groups. This test was chosen because it does not assume normality and is robust against outliers. To adjust for multiple testing and control the rate of type I errors, a false discovery rate (FDR) correction was performed using the Benjamini-Hochberg procedure.

Linear regression analysis was conducted to assess the correlation between metabolites and stroke outcomes. The regression models included sex, maternal diet, and their interaction terms to determine the influence of these factors on metabolite levels.

### Pathway and Enrichment Analysis

Pathway analysis was performed using MetaboAnalyst (version 5.0). The pathway analysis involved the use of the hypergeometric test and pathway topological analysis. The p-values were not Bonferroni corrected because the related hypotheses are not completely independent, but instead share underlying biological mechanisms or functional relationships. Enrichment analysis was performed to identify the over-representation of metabolites in specific pathways using Fisher’s exact test.

### Visualization

Heatmaps were generated to visualize the clustering of metabolites and samples. Pathway and enrichment analysis results were visualized using bubble plots and bar plots, respectively. The level of significance was set at p < 0.05 for all analyses.

## Acknowledgements

None

## Data statement

All relevant data can be found within the article and supplementary material

## Author contrition statement

Conceptualization: N.M.J.; Methodology: H.G., J.C.; Formal analysis: P.J., F.A., N.M.J; Writing-review & editing: F.A., M.T.M, P.J., H.G., N.M.J; Visualization: P.J., F.A., N.M.J; Project administration: N.M.J.; Funding acquisition: N.M.J.

## Diversity and inclusion statement

All authors were given opportunities to be involved in the study and writing up of the study.

